# A novel and highly divergent Canine Distemper Virus lineage causing distemper in ferrets in Australia

**DOI:** 10.1101/2021.11.03.467217

**Authors:** Ankita M. George, Michelle Wille, Jianning Wang, Keith Anderson, Shari Cohen, Jean Moselen, Leo Yi Yang Lee, Willy W. Suen, John Bingham, Antonia E Dalziel, Aeron C. Hurt, David T. Williams, Yi-Mo Deng, Ian G. Barr

## Abstract

Canine distemper virus (CDV) is a highly contagious systemic viral disease of dogs, that regularly spills-over into other animal species. Despite widespread vaccination, CDV remains endemic in many parts of the world. In this study we report an outbreak of distemper in ferrets in two independent research facilities in Australia. We found that disease severity varied, although most animals had mild to moderate disease signs. Histopathology results of animals with severe disease presented the typical profile of distemper pathology with multi-system virus replication. Through the development of a discriminatory PCR paired with full genome sequencing we revealed that the outbreak at both facilities was caused by a single, novel lineage of CDV. This lineage was highly divergent across the H gene, F signal peptide and full genome and had less than 93% similarity across the H gene to other described lineages, including the vaccine strain. Molecular analysis indicates that this strain belongs to a distinct lineage that diverged from other clades approximately 140 to 400 years ago, and appears to be unique to Australia. Given the differences in key viral proteins of this novel CDV strain, a review of the efficacy of the CDV vaccines currently in use in Australia is warranted to ensure maximum protection of dogs and other vulnerable species. In addition, enhanced surveillance to determine the prevalence of CDV in ferrets, dogs and other at-risk species in Australia would be useful to better understand the diversity of CDV in Australia.

**Importance:** Canine distemper virus (CDV) is highly contagious and while dogs are the main reservoir, it may spill over into a number of other animal species. In this study we report an outbreak of distemper in ferrets in two research facilities in Australia. Outcomes of pathology and histopathology suggest ferrets have widespread multi-systemic infection, consistent with previously reported distemper infections in ferrets and dogs. Critically, through sequencing and phylogenetic analysis, we revealed that the outbreak at both facilities was caused by a single, novel and highly divergent lineage of CDV. This virus had less than 93% nucleotide similarity to other described lineages and the vaccine strain. This manuscript adds considerably to the epidemiology, ecology and evolution of this virus, and is one of few reports of distemper in Australia in the literature.

## Introduction

Canine distemper virus (CDV) (species *Canine morbillivirus*), is a member of the genus *Morbillivirus*, in the family *Paramyxoviridae* (1) and causes endemic and widespread infectious disease in animals in many countries (2, 3). Canine Distemper Virus is a multi-host virus that infects animals from a wide taxonomic range, including canids, felids, mustelids, procyonids and phocids (3, 4).The main reservoir for this virus are dogs, but due to the broad host range (5), there is continual spill over between domesticated animals and wildlife (6). Disease signs include severe rash and both ocular and nasal discharge and over the course of infection animals often develop neurological signs including circling, hyperesthesia, seizures, cerebellar or vestibular disease, in the form of head tilting (7).

Severe systemic infection can occur in immunologically naïve animals (3, 8), with long periods of viral shedding, with infection in domestic dogs previously reported to last as long as 90 days (9). CDV is highly contagious in many species of animals (6), and resultant mortality rates vary in animal populations and across the species affected (8). For example, mortality has ranged from 23% in the Chinese giant panda (*Ailuropoda melanoleuca)* (10) to 100% in ferrets (*Mustela putorius furo*) (11). The origin and transmission of CDV infections, particularly in non-canidae species, such as ferrets, is poorly understood (9, 12, 13). Ferrets are a widely used experimental animal to study the pathogenesis and transmission of a variety of viral diseases including influenza, SARS-CoV-1 and −2, Ebola, rabies and a range of paramyxoviruses (14, 15), and are therefore an important animal to protect from unwanted, potentially fatal infections such as CDV.

Currently there are 17 distinct geographically-associated genetic lineages of CDV described globally (5, 16). On some continents there may be more than one lineage and for many, the evolutionary genetics have been well resolved (5, 17). However CDV has not been well documented in Australia (11), with only 48 reported cases in dogs and ferrets from 2006-2014, and limited information on the emergence and spread of CDV (11, 18, 19). This is further exacerbated by the total lack of publicly available genetic sequences from outbreaks in Australia.

There is a global vaccination strategy in place for the management and control of CDV with most countries, including Australia, generally utilizing a modified live attenuated vaccine based on the Onderstepoort strain. While this strategy is important in controlling this virus in dogs (8, 16, 20) there are a number of records where there is evidence of vaccine-induced CDV infections in both domesticated dogs and in wildlife through reversion of attenuation in the vaccine strain (6, 21). For example, CDV cases in wildlife due to the vaccine strain infection have been reported in South Africa and the UK (6, 22, 23). In Australia, routine vaccination is available for domesticated dogs, with vaccination normally given to puppies and other domestic animals at risk of contracting CDV, such as domesticated ferrets (11), using a live attenuated CDV. Despite a large-scale ongoing vaccination strategy in Australia, strains of CDV continue to circulate in animal populations (11). For example, in a study conducted from 2006-2014 in dogs and ferrets in Australia, it was reported that there were confirmed or suspected cases of CDV in five states/territories with most cases reported in New South Wales, including two out of three tested ferrets (11). However data pertaining to CDV epidemiology in Australia is limited, and the modes and rates of infection and mortality are not well understood.

In this study we report an outbreak of CDV in the ferret population in the Australian state of Victoria in 2019. CDV was detected in ferrets supplied to two independent research facilities from different breeders, indicating that this outbreak likely occurred across wide regions of the state. Herein we describe this CDV outbreak based on clinical signs and qRT-PCR CDV detection and describe the histopathology of severe cases. To better discriminate the dynamics of the outbreak we developed an assay to discriminate between a commercial CDV vaccine and the field strain. By full genome sequencing we also determined that the CDV in this outbreak originated from a novel lineage of CDV.

## Methods and Materials

### Ethics Statement

Work with ferrets at The Peter Doherty Institute for Infection and Immunity (hereafter Doherty Institute) was conducted with the approval of the Melbourne University Animal Ethics Committee (AEC# 1714278). Animals that had severe disease signs received veterinary intervention and were humanely euthanised in accordance with ethical guidelines. Experiments using ferrets at the Australian Centre for Disease Preparedness (ACDP) were approved by the ACDP Animal Ethics Committee (AEC# 1956). All work at the Doherty Institute and ACDP was performed in strict accordance with the Australian Government, National Health and Medical Research Council Australian code of practice for the care and use of animals for scientific purposes (24).

### Ferrets

Ferrets used for experimental purposes at the Doherty Institute and the ACDP were obtained from three different ferret breeders (2 supplied the Doherty Institute and 1 supplied ACDP) in the state of Victoria, Australia. The breeders are located at different locations outside of the Melbourne metropolitan region and provided ferrets at regular intervals to both facilities throughout 2019.

### CDV Vaccines

The Protech C3 vaccine (Boehringer Ingelheim, Australia), comprising a live-attenuated CDV was administered at both the Doherty Institute and ACDP to control the respective CDV outbreaks. The dose administered varied between institutions: at the Doherty Institute 0.2ml was administered to each ferret and at the ACDP 0.25ml was administered, regardless of age or weight. At both facilities two doses were administered, however at the Doherty Institute doses were 2 weeks apart and at the ACDP doses were administered 4 weeks apart. The Protech C3 vaccine has been approved for dogs in Australia and the standard dose administered to dogs is 1ml.

### Histology and Immunohistochemistry

Tissues collected for histology processing at ACDP were fixed in 10% neutral buffered formalin, processed and embedded in paraffin using standard procedures, sectioned at 4 µm, and stained with hematoxylin and eosin (H&E). For immunohistochemistry (IHC), paraffin-embedded tissue sections were quenched for 10 min in aqueous 10% hydrogen peroxide. Antigen retrieval was performed by using the Agilent PT Link (Dako, Agilent, Vic, Australia) module for 30 min at 97°C in pH 9 antigen retrieval solution. A mouse monoclonal antibody targeting the nucleocapsid protein of CDV (CDV-NP, VMRD, WA, USA) was used at a dilution of 1:2000 (60 min incubation), sections were incubated with Mouse linker (Dako), then visualised using an Envision Flex horseradish peroxidase (HRP)-secondary antibody (DM822, Dako) for 20 minutes (goat anti-rabbit and anti-mouse immunoglobulins), followed by chromogen aminoethyl carbazole (AEC), and then counterstained with Lillie-Mayer’s Haematoxylin and Scotts Tap Water. Sections were digitised using a Pannoramic Scan II (3DHISTECH Ltd, Budapest, Hungary) whole slide imager before photomicrographs were taken using the image capture function of the CaseViewer software (3DHISTECH Ltd).

### Identification of ferrets with CDV by Real-Time PCR

Two different PCR approaches were used across the two research facilities. At the Doherty Institute a commercial CDV quantitative real-time reverse-transcriptase PCR (qRT-PCR) was initially employed followed by the development of a discriminatory assay. All ferrets received into the facility were tested. At ACDP, a pan-morbillivirus RT-PCR was applied for detection of CDV from ferrets with clinical symptoms (25).

At the Doherty Institute nasal wash samples were collected from lightly sedated ferrets. Ferrets were sedated by intramuscular injection of a combination of ketamine (12.5mg/kg, Troy Laboratories) and xylazine (2.5mg/kg, Troy Laboratories). We instilled 1 mL of sterile PBS into one nostril and allowing the liquid to flow out of the other nostril into a collection tube. Nasal wash samples were immediately stored at −80°C until RNA extraction. RNA was extracted from 140µl of nasal wash sample using the QIAamp Viral RNA Mini kit (QIAgen, Australia), according to the manufacturer’s instructions. RNA was also extracted from the Protech C3 vaccine (Boehringer Ingelheim, Australia) following reconstitution according to manufacturer’s instructions.

Primers were developed for a number of purposes (Table 1). First, to develop a discriminatory PCR assay to distinguish between the outbreak virus and the vaccine virus, second to amplify genes of interest (NP, H, and F) and finally for whole genome sequencing (WGS) (Table 1). At the Doherty Institute, primers designed in this study were created from conserved regions of similar sequences to the wild type strain of CDV found in Victorian ferrets (percentage identity ∼92%) retrieved from GenBank using (Basic Local Alignment Tool) Blastn. Primers were created with both Primer-BLAST (NCBI), and Geneious R10 (Biomatters, Auckland, New Zealand).

**Table 1.**
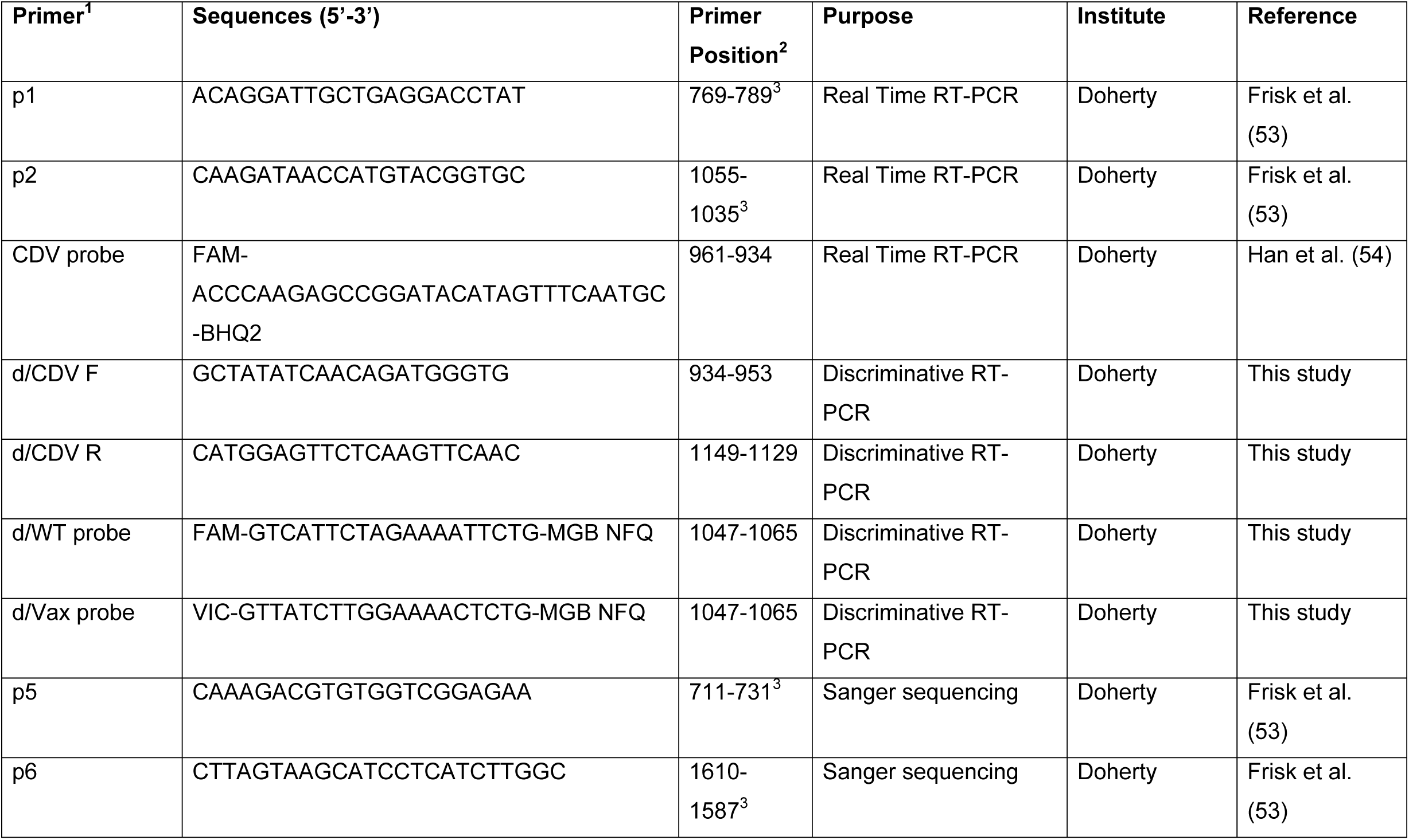
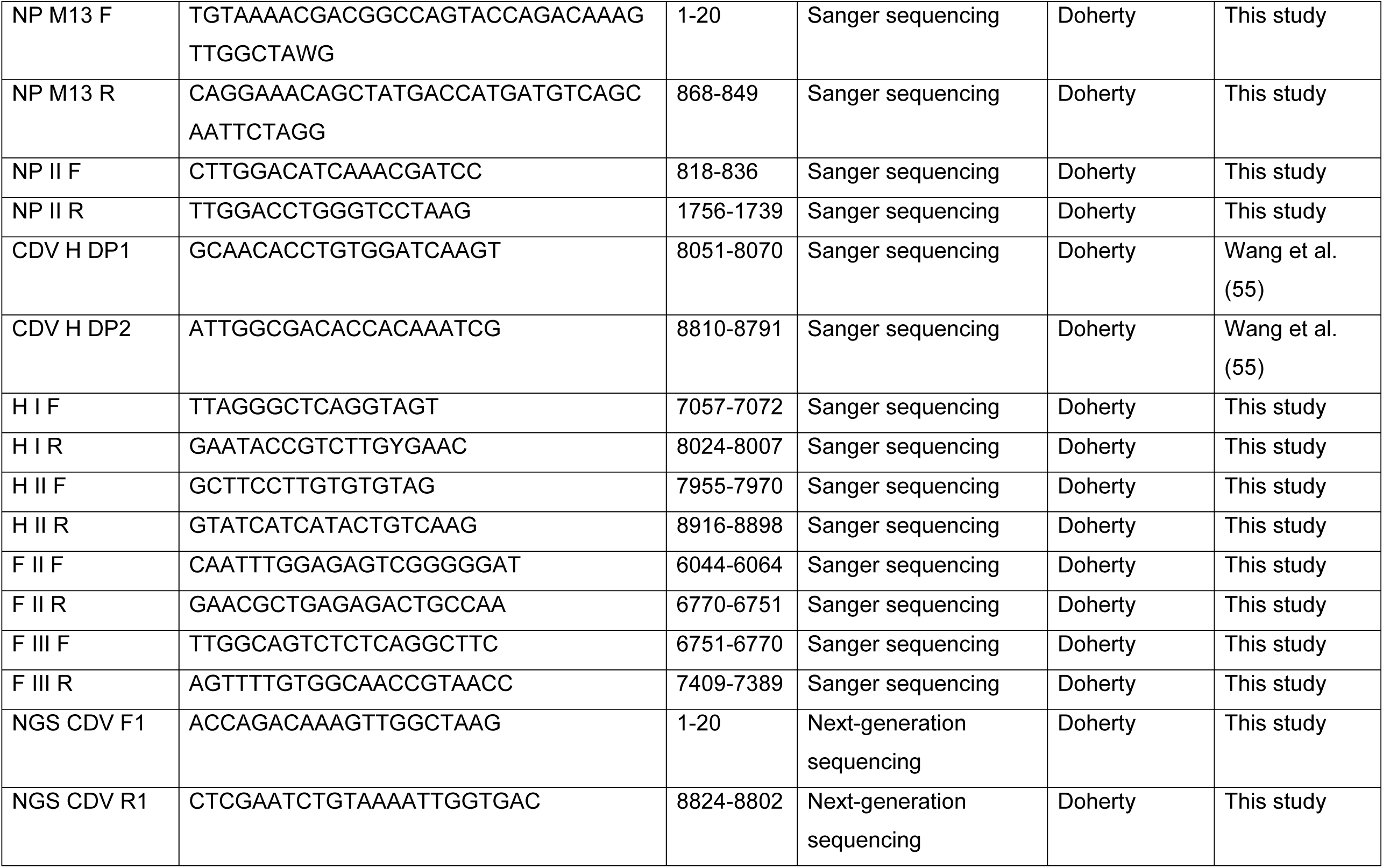
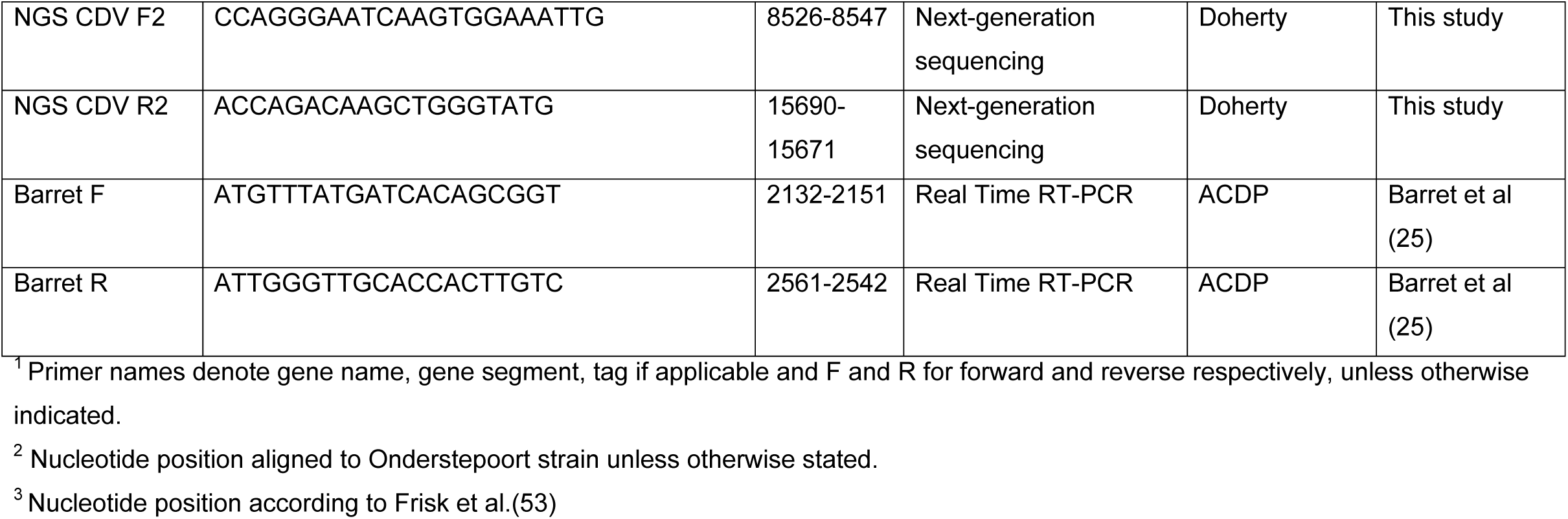
Primers and probes used at the Doherty Institute and ACDP for the detection of CDV in ferrets

qRT-PCR was performed using the SensiFAST Probe Lo-ROX One-Step kit (Bioline Meridian, Australia) with a reaction volume of 20µl. The reaction mix contained 4ul RNA, 40µM of each primer and 10µm of the associated probe. Thermocycling conditions comprised a reverse transcriptase step at 45°C for 10 minutes, followed by a denaturation step at 95°C for 2 minutes and 40 cycles of 95°C for 5 seconds and 60°C for 15 sec. The assay designed here was validated against a commercially available CDV assay the Canine Distemper Virus Detection qPCR (Genesig, United Kingdom).

To discriminate between the outbreak virus and the vaccine strain a discriminative qRT-PCR was developed using a two-probe assay (Table 1) with the SensiFAST Probe Lo-ROX One-Step kit (Meridian Biosciences). The reaction mix contained 10µM of each probe, 40µM of each primer with a final reaction volume of 20µl and the same thermocycling conditions were used as specified above.

At ACDP, both fresh and formalin-fixed-paraffin-embedded (FFPE) tissues including brains and nasal swabs from ferrets with clinical symptoms were used for RNA extraction and PCR testing. Total RNA was extracted from fresh or FFPE tissue samples. For fresh tissue samples, fifty microlitre of supernatant of 10% tissue homogenates from relevant ferrets was utilised by using the MagMax 96 Viral RNA Kit (ThermoFisher Scientific) in a MagMAX Express Magnetic Particle Processor (ThermoFisher Scientific) following manufacturer’s instructions. For FFPE samples, RNA was extracted from 4 uM sections of formalin-fixed, paraffin-embedded brain/lung tissue, using RNeasy FFPE Kit (QIAGEN), following the manufacturer’s instructions. The RNA was used for RT-PCR and Next-generation Sequencing (NGS) analysis.

A pan-morbillivirus RT-PCR, targeting the phosphate (P) gene of morbillivirus, was utilised for detection of CDV from ferrets with clinical symptoms (25) (Table 1). The RT-PCR assays were conducted by using SuperScript III One-Step RT-PCR System with Platinum Taq DNA polymerase (Invitrogen, Carlsbad, CA, USA). Amplifications were performed in an Eppendorf Master Cycler Model 5345 (Eppendorf, Hamburg, Germany). Each 25 uL reaction contained 12.5 uL 2X reaction mix, 1 uL Superscript III RT/Platinum Taq mix, 0.9 µM (final concentration) of each primer and 5 mL RNA template. The RT-PCR cycling conditions were as follows: one cycle of 48°C for 30 min for cDNA synthesis; one cycle of 94°C for 2 min for denaturing; followed by 45 amplification cycles of 94°C for 30 s, 50°C for 60 s, 68°C for 60 s and a final extension cycle of 68°C for 7 min. Amplified PCR products (approx. 429 bp for P gene) were gel purified using QIAquick gel extraction kit (QIAGEN, Hilden, Germany) and sequenced using BigDye Terminator v3.1 Cycling Sequencing Kit (Applied Biosystems, Foster City, CA, USA) for confirmation.

Prevalence of CDV at the Doherty Institute was calculated using the *bioconf()* function in the Hmisc package *(26)* and prevalence differences between the field strain and vaccine strain at the Doherty Institute ferret facility were compared using a Chi-squared test.

Differences in prevalence over time were compared using a generalized linear model with a binomial response and the statistically significant model was plotted using the *ggplot2* package (27). Analyses were conducted using R 4.0.2. integrated into RStudio 1.3.1073.

### Sanger Sequencing and Next-Generation

Initial sequencing was done at the Doherty Institute with Sanger sequencing. RNA that was stored at −80°C and had undergone limited freeze thaw cycles was used for RT-PCR using the MyTaq One step RT-PCR kit (Meridian Biosciences) and primers designed for Sanger Sequencing (Table 1). Product sizes were confirmed using the e-Gel 2% agarose (GP) (Invitrogen, USA) followed by purification using the Exo-SAP-IT PCR product clean-up reagent (Thermofisher, Australia) according to manufacturer instructions. The purified template then underwent sequencing using the BigDye Terminator v3.1 Cycle sequencing kit (Thermofisher). The primers used for this step were 10% the concentration of the same primers used to generate PCR products, unless the primer had a M13 tail, in which case a M13 F(5’-TGTAAAACGACGGCCAGT-3’) and R (5’-CAGGAAACAGCTATGACC3’) was used. Sequencing was performed on a 3500XL Genetic Analyser (Applied Biosystems). Results were analysed using Lasergene 13 (DNASTAR, Madison, WI, USA).

For subsequent next-generation sequencing (NGS) at the Doherty Institute, RNA was extracted from stored (−-80°C) nasal washes as described previously. Briefly, RT-PCR products were created using the Superscript IV One Step RT-PCR system (Thermofisher) and primers designed based on the results of Sanger Sequencing (Table 1). PCR product concentration was determined using a 4200 TapeStation System (Agilent Technologies, USA) and Genomic DNA Screentape (Agilent Technologies). The concentration of each sample was normalised to 6.6ng/µl and 30µl of DNA was used for Nextera DNA Flex Library Prep (Illumina, USA), performed according to manufacturer’s instructions. Sample libraries were pooled and diluted to 200pm and 100µl of this diluted sample was loaded onto a flow cell for NGS using an iSeq 100 (Illumina). Sequences were assembled using reference mapping with paired ends in Geneious R10 (Biomatters) and Bowtie2 (28) was used for end to end alignment.

RNA extracted from both fresh and FFPE tissue samples from ferrets tested positive by pan-morbillivirus RT-PCR was used for NGS at ACDP. Both field strains sequenced comprised of FFPE samples, and all but one vaccine strains sequenced were from fresh samples.

At ACDP, the TruSeq RNA Library Prep Kit (Illumina, USA), following manufacturer’s instructions was used for library construction, using an RNA concentration of 5 ng/µl. The libraries were normalised and pooled at equimolar ratios (final concentration of 12.5 pm each). The library pool was then loaded into flow cell of MiSeq Reagent Kit V2 (2 × 150 cycles) and sequenced in a MiSeq platform (Illumina), according to the manufacturer’s instructions. The NGS sequence data was analysed using CLC Genomic Workbench 20 (Qiagen) using standard parameters. The raw reads were quality-trimmed and viruses were assembled using a combination of read mapping (stand-alone read mapping, CLC Workbench algorithm) and *de novo* assembly (scaffolding, CLC Workbech algorithm) to generate consensus sequences.

All CDV sequences have been deposited in GenBank (Accession number: XXXX-XXXX). Sequences derived from the Protech C3 vaccine have been deposited with permission from the manufacturer (Boehringer Ingelheim).

### Phylogenetic Analysis

Sequences were assembled in SeqMan Pro 13 (DNASTAR). Assembled sequences were compared through the using BLASTn (NCBI) to other available sequences in the non-redundant nucleotide (nr) database (29).

To reconstruct the evolutionary relationships, all CDV sequences available in GenBank were downloaded and all geographic and host species origins were cross referenced with publications if they were not available in the GenBank record. Full genomes, coding complete H gene or the 135bp F signal peptide were aligned using the MAFFT algorithm (30) executed in Geneious R10 (Biomatters). Maximum likelihood trees were estimated using PhyML (31) and implementing the appropriate nucleotide substitution model. Lineage information and appropriate rooting was mined from Duque-Valencia et al.(5).

A time-structured phylogeny was constructed using the same backbone sequences and parameters as previously published (17). Briefly, select sequences were downloaded and the H gene was aligned using MAFFT and a maximum likelihood tree was constructed using PhyML. The tree was imported into TempESt v1.5.3 (32) to ensure clock-like behaviour in the data. Using BEAST 1.10.4 (33), the data were analysed using a strict clock, the HKY+G+I substitution model and a constant size coalescent model. The analysis was run for 100,000,000 generation and convergence was assessed using Tracer V1.6. A maximum clade credibility tree was generated using TreeAnnotator v1.8 and visualized in FigTree v1.4

## Results

### The Doherty Institute outbreak

Starting in April 2019, ferrets with clinical disease consistent with CDV were observed in the Doherty Institute animal facilities during routine animal husbandry. Nasal wash samples were collected from animals with clinical signs (as described below) and tested for CDV using the Canine Distemper Virus Detection qPCR (Genesig). In response, the Doherty Institute animal facility introduced routine vaccination of ferrets prior to their arrival at the Institute, starting in mid April of 2019, 10 days following the confirmation of the index case of CDV (Fig 1).

**Figure 1.**
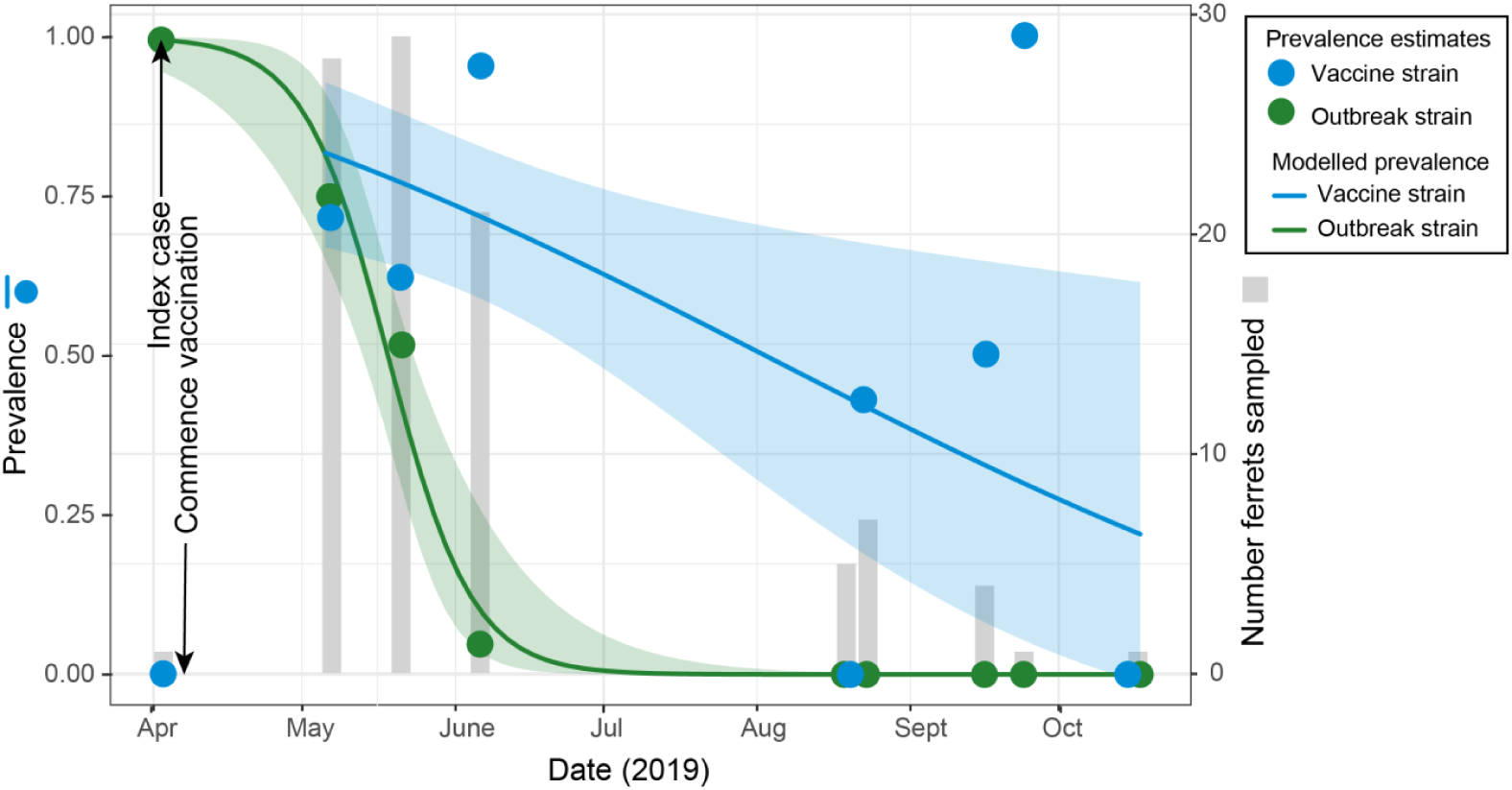
Time series of vaccine strain and field strain CDV prevalence in the Doherty Institute ferret facility. Grey bars indicate when, and the number of samples tested using the discriminatory PCR developed here and tick marks are on the secondary Y axis. Points represent the individual point estimates for the vaccine strain and the field strain. Lines correspond to modelled prevalence using a generalized linear model and shaded areas represent the 95% confidence interval of the model. We were unable to generate a corresponding outbreak figure for ACDP as they did not use a discriminatory PCR.

Between April 2019 and December 2019, all 179 animals that entered the Doherty Institute Biological Resource Facility (BRF) were tested for CDV following arrival. Of these animals, 115 were positive (64%) for CDV by RT-PCR. Using the discriminative assay developed here, we tested 103 of these 115 CDV positive samples and detected two different strains of CDV – a field strain causing the initial outbreak and the strain used in the Protech C3 vaccine. Of the 103 animals screened with the discriminative assay, 100 were vaccinated prior to sampling (approximately 3 weeks prior to nasal wash sampling). Overall, there was a statistically significant difference in the proportion of detections due to the field strain (39%) and vaccine strain (56%) (*X*^2^=12.921, df=1, p=0.003) (Fig 1). In 20 ferrets, both the vaccine and field strain of the virus were detected. Despite finding the vaccine strain in all sampling time periods following the routine introduction of CDV vaccination and having a higher overall prevalence, we found that the CDV outbreak caused by the field strain persisted for less than two months (Fig 1). Following the index cases on 3 April 2019, the field strain was detected in approximately 70% of animals on 7 May, 50% on 21 May, but was no longer detected in ferrets from June 2019. This is in contrast to the vaccine strain, which was consistently detected during the seven months following CDV vaccine implementation on 3 April (Fig 1). There was no statistically significant difference in disease outcomes (*i*.*e*. requiring euthanasia under the ethical guidelines) when comparing ferrets infected with the field strain and the vaccine strain; 3 ferrets positive for the vaccine strain were euthanised and 3 ferrets infected with the field strain were euthanised. An additional ferret that was negative for CDV was euthanised, and three additional ferrets that were not tested for CDV were euthanised (Table 2).

**Table 2.** Number of ferrets infected with CDV in the Doherty Institute.

| Ferrets | qPCR<br>Positive<sup>1</sup> | Positive for field<br>strain <sup>2</sup> | Positive for<br>Vaccine strain<sup>2</sup> | Negative CDV<br>result<sup>1</sup> |
| --- | --- | --- | --- | --- |
| **Tested** | 115 | 41 | 65 | 64 |
| **Euthanised** | 6 | 3 | 3 | 1 |

1. Based on either the qPCR assay from relevant literature (53, 54) and/or the commercial Canine Distemper Virus Detection qPCR
2. Based on the discriminatory PCR developed in this study

### ACDP Outbreak

Coinciding with the first detection at the Doherty Institute animal facility (April 2019), ferrets at the animal holding facility for ACDP (CSIRO Werribee Animal Health Facility) began to experience clinical signs consistent with CDV infection (as described below). Unlike the Doherty Institute, no qPCR or discriminatory RT-PCR was leveraged to identify CDV positive ferrets. PCR was only used retrospectively. Rather, in response to clinical signs, quarantine measures were initially introduced to separate ferrets with clinical signs from unaffected ones. All ferrets with clinical signs and animals sharing cages regardless of clinical signs were euthanised to control the outbreak. A detailed breakdown of infection status and euthanasia from the ACDP outbreak are not available. Vaccination was introduced using a quarter of the standard dose of the Protech C3 vaccine. The initiation of vaccination coincided with the final detections of CDV in the facility; within two months of the initial vaccination, the CDV outbreak at ACDP was controlled.

### Clinical Signs

Clinical signs of CDV observed in ferrets from both the Doherty Institute and ACDP included (Fig 2A) inguinal dermatitis, with rough, discoloured patches of skin appearing on the abdomen, (Fig 2B) hyperkeratosis of the footpads, also known as hardpad, (Fig 2C) dermatitis on the chin and mouth, with scaly patches forming, and/or ocular signs including uncontrolled eye twitching with mucopurulent ocular and nasal discharge, resulting in visible crusting around the eyes and nose (Fig 2). At the Doherty Institute, visible signs were mostly classified as mild, with the animals remaining active. However, 10 animals (including both animals that were not tested by qPCR [n=3] and those that were tested for CDV by qPCR [n=7] displayed more severe signs (Fig. 2C-D) and required immediate euthanasia in accordance with ethical requirements. Animals classified as suffering from severe disease also showed signs of decreased activity, lethargy and loss of appetite. Any one of these clinical signs as well as the appearance of lethargy, pyrexia, anorexia, or weight loss were determined to be significant enough to require veterinary intervention. Due to the implementation of a euthanasia strategy to contain the outbreak, a breakdown of disease severity was not available from ACDP. Necropsies were performed on all euthanised animals at the Doherty Institute and on selected cases at ACDP. Tissues taken at the Doherty Institute were sent to an external pathology service, however detailed results were not available. Tissues collected at ACDP were processed for histopathology assessment at ACDP, as described below.

**Figure 2.**
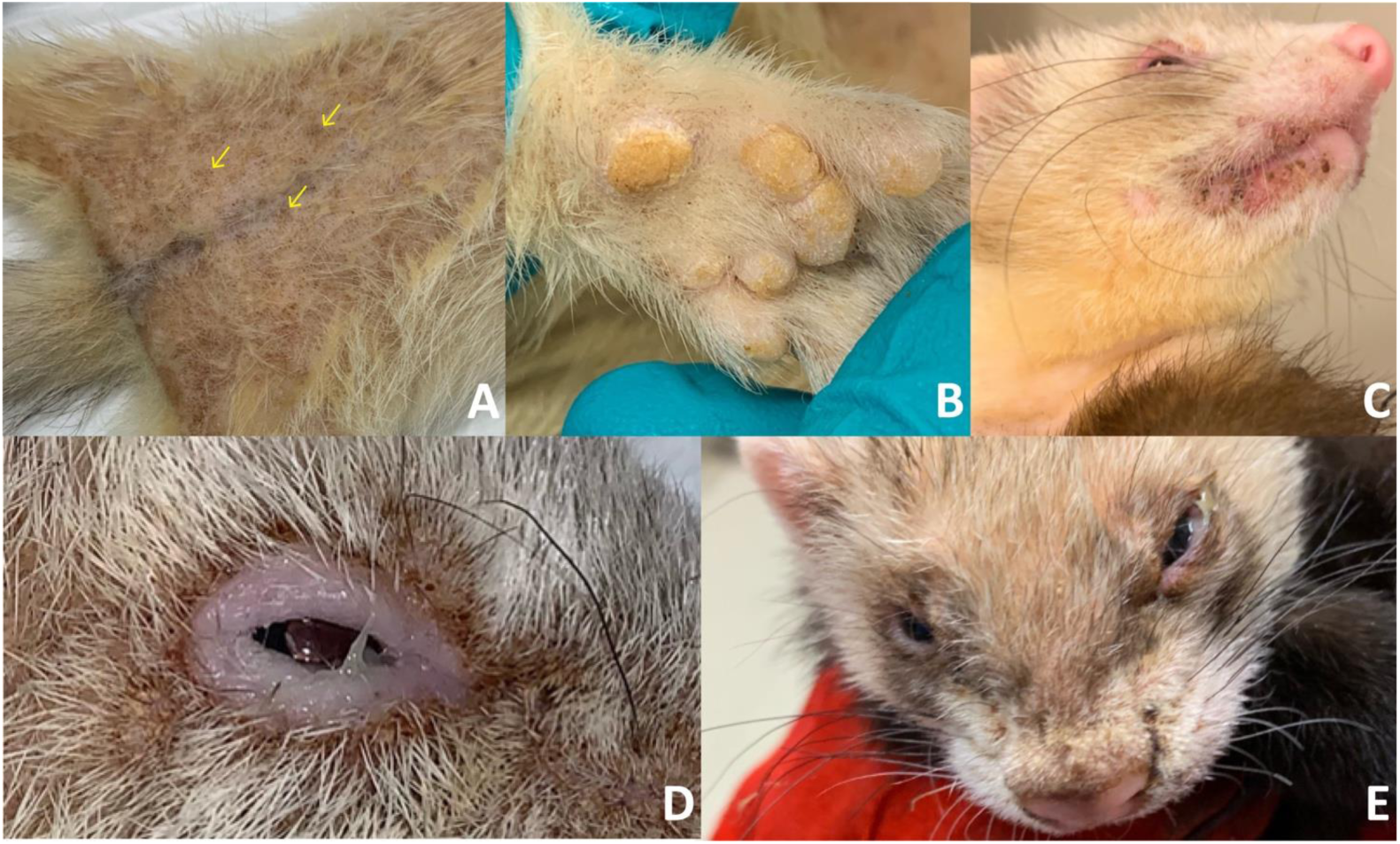
Clinical signs of animals infected with CDV. Animals that developed visible signs of CDV had (A) mild abdominal rashes (as indicated by arrows) and (B) crusting of footpads. One animal developed (C) crusting around the mouth, however the majority of visible symptoms on the face consisted of crusting around the (D) eyes and (E) nose.

### Histopathology and CDV immunohistochemistry

Four ferrets with distemper-like clinical signs and one asymptomatic ferret from the ACDP outbreak were submitted for histopathological analysis. Three of the symptomatic ferrets were submitted early in the outbreak, another symptomatic ferret was submitted days after vaccination was introduced, along with an asymptomatic penmate. In these ferrets, necrosis and sloughing of the bronchiolar epithelium were observed in the lung. This was often accompanied by notable diffuse epithelial hyperplasia (Fig 3A) and occasionally with a lymphohistiocytic infiltrate. In some of the affected areas, frequent round eosinophilic inclusion bodies were detected in the cytoplasm and nucleus of bronchial/bronchiolar epithelial cells (Fig 3A). Occasional multinucleated syncytial cells were found lining alveoli. Similar syncytial cells and intracytoplasmic/intranuclear inclusion bodies were identified in hyperplastic parts of the renal pelvic and urinary bladder urothelium (Fig 3B). The skin over the nasal planum was also hyperplastic with prominent parakeratotic hyperkeratosis emanating most notably from hair follicles. Associated suppurative inflammation was also occasionally observed in the subjacent dermis.

**Figure 3.**
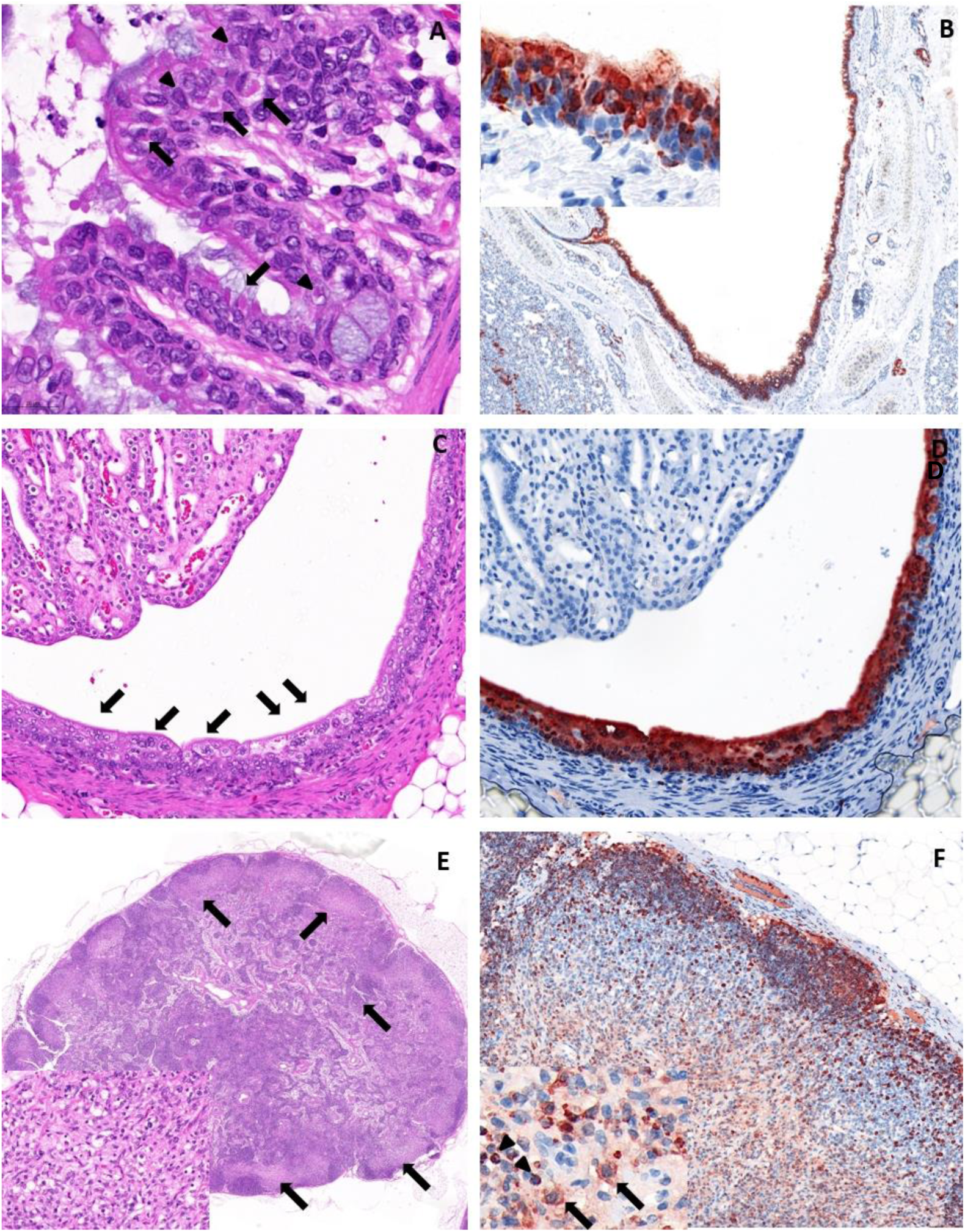
Histopathology and immunohistochemistry (IHC) of ferrets from the CSIRO ACDP outbreak. (A) H&E stain of the lung showing multiple eosinophilic intracytoplasmic (arrows) and intranuclear (arrowheads) inclusion bodies within bronchiolar epithelial cells. 40x magnification. (B) CDV IHC of the lung with diffuse intense cytoplasmic labelling in the bronchial epithelium. 5x magnification. Inset shows the homogeneous to coarse granular and globular cytoplasmic labelling in the bronchial epithelium. 40x magnification. (C) H&E stain of the renal pelvis with multiple syncytial cells in the urothelium (arrows). 10x magnification. (D) CDV IHC of the renal pelvis, corresponding to panel C, with diffuse intense cytoplasmic labelling in the urothelium. 10x magnification. (E) H&E stained section of a lymph node with severe coalescing areas of lymphoid depletion in the cortex. These areas were replaced with patches of fibrin, oedema, histocytes and early fibroplasia (arrows and inset). Inset in panel E. corresponds to an area of lymphoid depletion as indicated by the arrows in panel E. 20x magnification. (F) CDV IHC of the lymph node corresponding to panel E illustrating antigen positive round cells were identified in the depleted cortex. 10x magnification. Inset shows antigen positive lymphocytes (arrowhead) and histiocytes (arrows) in an area of lymphocyte-depleted cortex. 40x magnification.

In addition to above lesions, multifocal to coalescing patches of the lymph node cortex were depleted in lymphocytes (Fig 3C). In the more severely affected lymph nodes, these lymphocyte-depleted areas in the cortex were replaced by oedema, fibrin, histiocytes and fibroplasia (Fig 3C). Taken together, the histopathologic features described above were consistent with pathology typically associated with CDV infection in ferrets.

Immunohistochemistry targeting the nucleocapsid protein of CDV showed that viral replication was widespread and intense, affecting many organ systems, even in animals with minimal histopathologic changes. Antigen was detected in the epithelium of the respiratory tract (Fig 3D), renal pelvis (Fig 3E), skin of the nasal planum, biliary tract, alimentary tract, female reproductive tract, lacrimal gland and salivary gland. High viral antigen burden was also observed in lymph nodes and spleen. Round cells with morphology consistent with histiocytes and lymphocytes were the main targets in these lymphoid organs (Fig 3F). In the brain, viral antigen was detected in only few mononuclear cells infiltrating the choroid plexus. Rare antigen positive round cells were also identified scattered in the olfactory bulb. This widespread multi-systemic infection was consistent with previously reported distemper infections in ferrets and dogs (34, 35).

### Sequence analysis and evolutionary genetics

We sequenced 3 complete genomes corresponding to the field strain of CDV, in addition to 8 complete genomes of the CDV vaccine strain from ferrets. Both the field and vaccine strain viral genomes sequenced at the Doherty Institute were recovered from nasal wash samples. The field strain viral genomes from ACDP were sequenced from FFPE brains. Most vaccine strain viral genomes from ACDP were sequenced from fresh tissue samples (nasal swabs and brain) and only one was from FFPE brain. Analysis of the three field strain genomes sequenced across the two facilities showed only limited genetic diversity, with 99.78% nucleotide similarity in the H gene, 98.1% in the Fsp region and 99.5% across the full genome. This strongly suggests that the outbreaks in both facilities were due to the same virus strain; the most parsimonious explanation is that the outbreak was initiated in the breeding facilities supplying both the Doherty Institute and ACDP. As expected, there was limited diversity in the virus sequences from animals infected with the vaccine strains that were sequenced (n=7) with 99.83% similarity in the H gene, 98.5% in the Fsp region and 99.8% across the full genome. The genome sequences corresponding to the vaccine strain and the Onderstepoort strain shared 99% nucleotide similarity in the H gene, 98.5% in the Fsp region and 99.4% across the full genome.

Analysis of the H (Fig 4) and Fsp (Fig 5A) genes of the field strain viral sequences, regions of the CDV genome currently used for lineage discrimination (17), clearly demonstrated the divergent nature of the field strain. Indeed, based upon these genes and the full genome sequences (Fig 5B), and the current approach for lineage designation (*e*.*g*. (17), the field strain should be designated as a novel lineage. Critically, the field strain is highly divergent from both the Protech C3 vaccine itself and from virus sequences from ferrets vaccinated with this attenuated strain. More specifically, the H gene shared less than 93% nucleotide identity with all existing lineages of CDV, including the Protech C3 vaccine sequence [92.5% nucleotide similarity, 91.8% amino acid similarity] which we also sequenced in this study (Fig 4). Phylogenetically, the outbreak virus belonged to the lineage comprising “Vaccines (both the Protech C3 and the Onderstepoort strain which is used in most attenuated vaccines globally), Asia-3 and North America −1”, as defined by (5) (Fig 4). Furthermore, this sequence is highly divergent from other CDV sequences from mustelids, globally, with the exception of 4 sequences from China which fall into the “Asia-3” lineage (Fig 4). Other mustelid sequences fell into an array of lineages, although the vast majority belonged to the “Asia-1” lineage (Fig 4). As outlined in previous studies, the H gene phylogeny is dictated largely by geography, and we speculate that the field strain reported here is representative of CDV currently circulating in Australia, however, as there are no available sequences in GenBank with which to compare, this cannot be confirmed.

**Figure 4.**
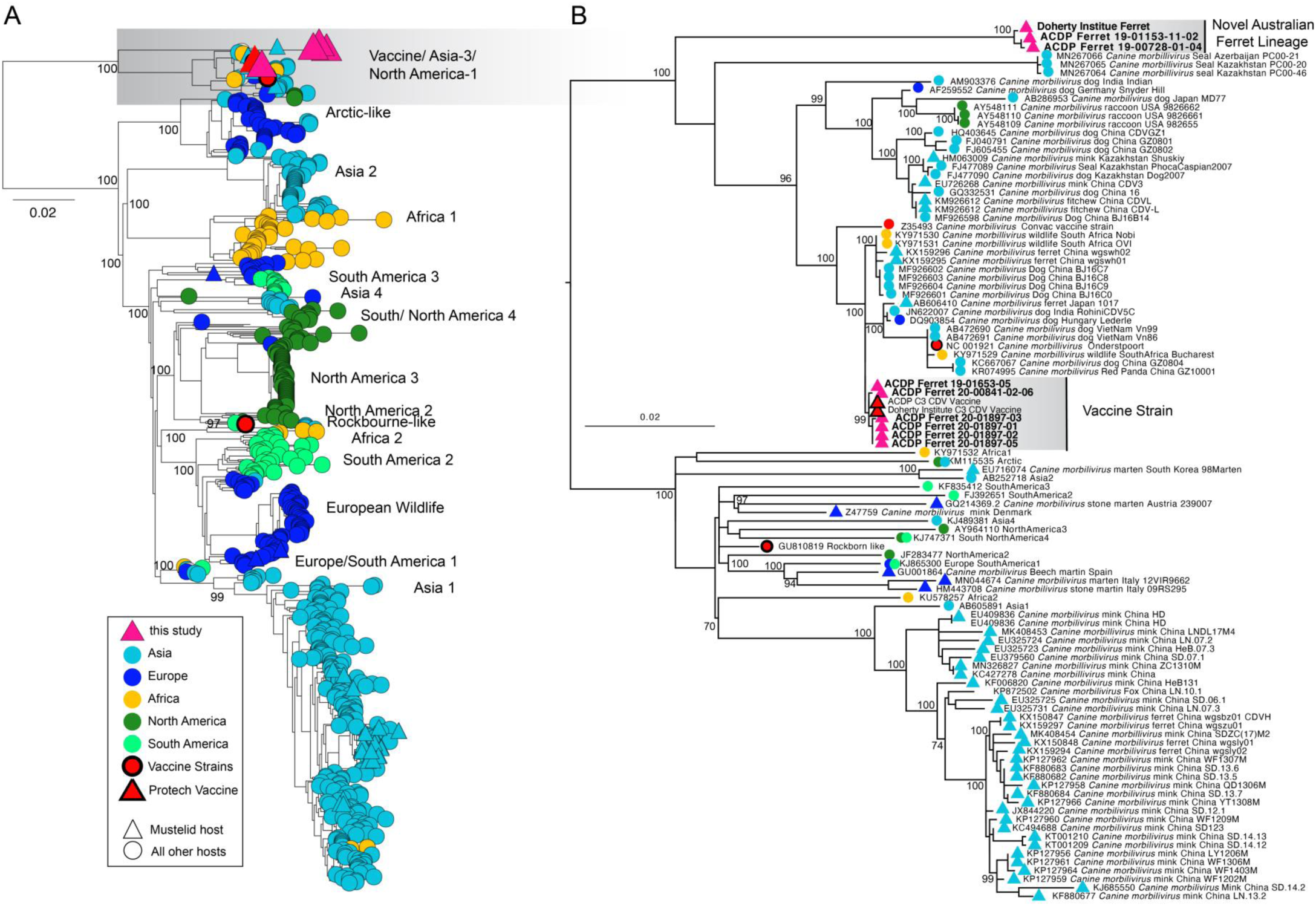
Phylogenetic analysis of the H gene of CDV. (A) Tree containing H gene of all CDV sequences available in GenBank. Tips are coloured by geographic region. Red shapes with a thick black border corresponds to a vaccine strain. Shapes indicate host, with all mustelid hosts indicated by a triangle. (B) Expansion of the “Vaccine/Asia-3/North America-1” lineage which is indicated by a grey box in A, in addition to reference sequences for main lineages and all sequences from mustelids. Sequences generated in this study are presented in a grey and sequences from ferrets are in bold text. Both the Doherty Institute and ACDP generated a full genome of the Protech Vaccine Strain, indicated by a red triangle in the “Vaccine strain” clade. Scale bar indicates the number of substitutions per site. Support values are presented for major nodes.

**Figure 5:**
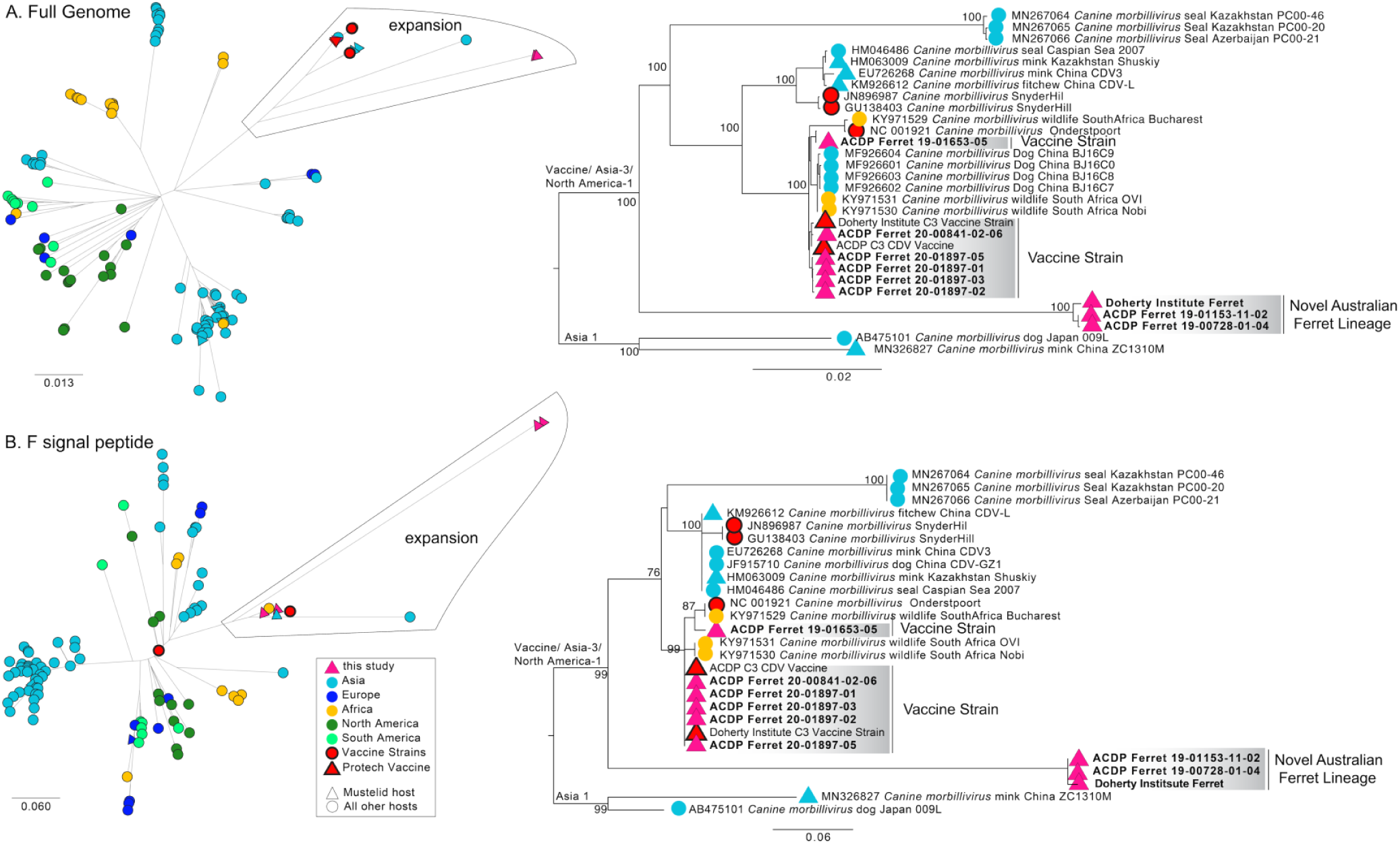
Phylogenetic analysis of (A) the full genome and (B) the F gene signal peptide. Trees containing all sequences in Genbank are unrooted, and the subtree that is expanded is denoted. Tips are coloured by geographic region or vaccine strain. Shapes indicate host. Sequences generated in this study are presented in a grey box in B. Scale bar indicates the number of substitutions per site. Support values are presented for major nodes.

We found similar patterns in the Fsp gene and full genome analyses (Fig 4). Phylogenetic analysis consistently demonstrated the field strain as being highly divergent (Fig 5), with the closest lineage being the “Vaccine/Asia-3/North America-1” clade and with high divergence from other mustelid CDV sequences. The Fsp region analysis of the novel Australian ferret lineage showed high divergence from other sequences, with only 73.88% of the nucleotide identity shared by both the vaccine and the Onderstepoort strain (Fig. 5B) compared to analysis of the full genome which shares approximately 91% identity with other reported strains (Fig. 5A).

Utilizing the framework presented by Duque-Valencia et al (2019) we aimed to estimate the divergence of the CDV viruses presented in this study (Fig 6). As we used the same BEAST parameters, including the use of a strict molecular clock, we were unable to include viruses from the “Vaccine clade” as these viruses have been passaged in cells and don’t have the same evolutionary rate as viruses infecting animals. Using this approach, we estimated the date that the field strain described here has branched from other clades between 1623.96 – 1878.9 (95% highest posterior density [HPD]) (Fig 6). It is also worth noting that with the addition of the sequences we generated in this study, tMRCA of the entire tree is significantly older than that presented in Duque-Valencia et al. (2019) but aligns more closely with Jo et al. (2019). While this analysis is useful in demonstrating the substantial period of time since these sequences have diverged from the closest lineage, it is important to take the specific dates merely as a guide. By also running this tree with an uncorrelated relaxed lognormal clock we were able to incorporate the vaccine strain, however this approach moves tMRCA much further back in time. This approach confirms that the divergence of CDV strains causing the outbreak described in this study occurs before the bifurcation of the “vaccine” clade from the “Asia-3” and “North America-1” clades, which is illustrated in all the maximum likelihood trees presented (Fig 4, 5).

**Figure 6.**
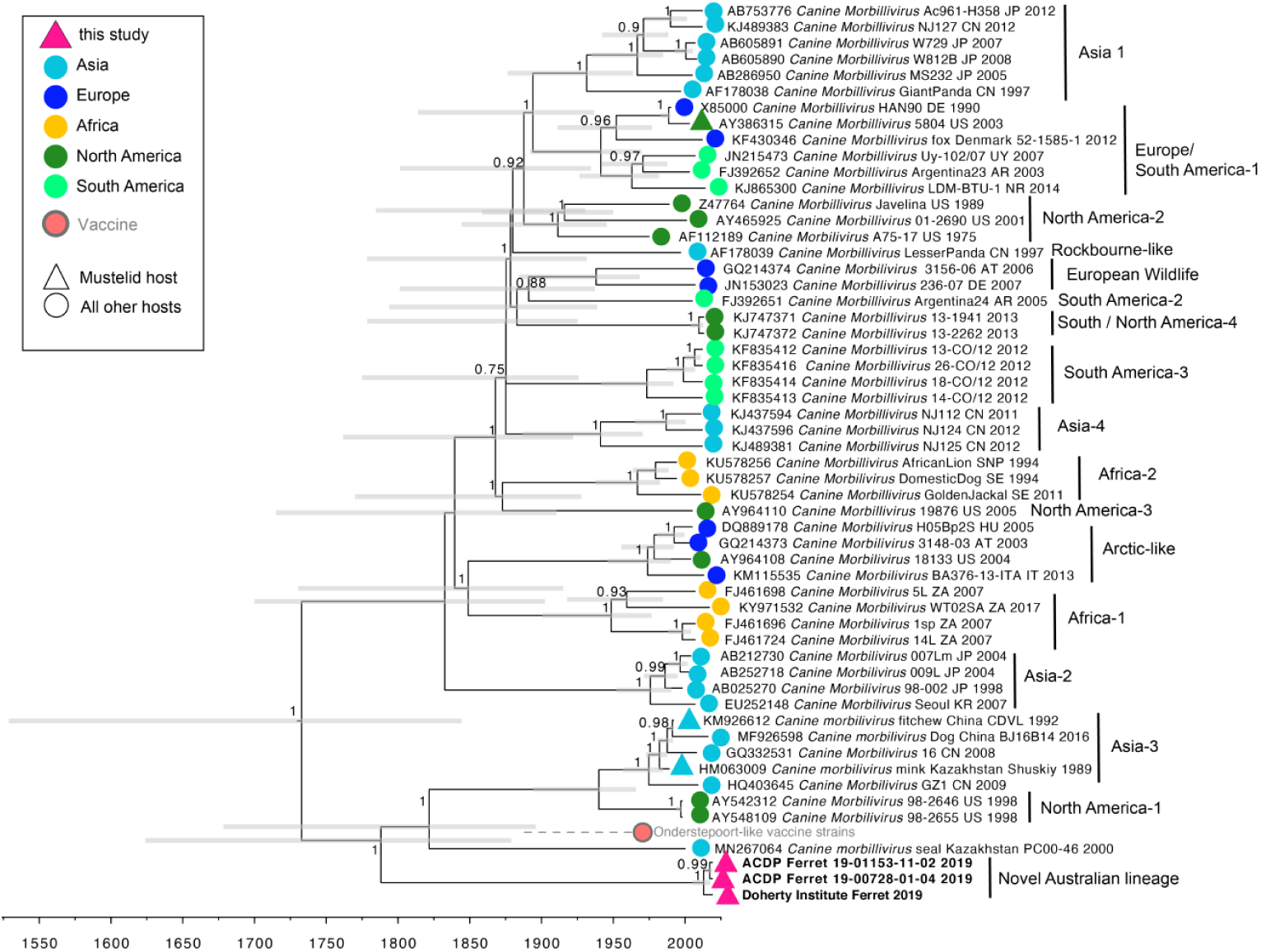
Time structured phylogeny of the H gene. Reference sequences are those presented in Duque-Valencia et al (2019). We did not include any vaccine strains, including the vaccine strains from ferrets sequenced in this study because repeated passage of vaccine strains in laboratory settings does not reflect natural evolution. However we have included the tentative position of the vaccine clade for clarity only – the outgroup to Asia-3 and North America-1. This phylogenetic position is based upon maximum likelihood estimation in this study and Duque-Valencia et al (2019). Field strain sequences generated in this study are in bold. Scale bar represents time in years. Node labels correspond to posterior probabilities of each node. Grey bars comprise the 95% highest posterior density of the date estimate.

## Discussion

The outbreak of distemper in ferrets in Victoria, Australia in 2019 described here represents one of the few outbreak investigations of CDV reported in Australia. Further, it is the first to describe and analyse the evolutionary genetics of CDV in Australia. Despite the new data presented here, the epidemiology, ecology and evolution of CDV in Australia remains entirely unknown, including the source and reservoir of the CDV lineage that infected the ferrets reported in this study.

The first documented CDV outbreak in Australia occurred in 1983 and involved over 1000 cases across 3 states and was associated with racing Greyhounds (19). Since then, outbreaks have been limited and with only a few reports. There are three contemporary reports: one recorded outbreak comprising 16 cases in 2004-05 in unvaccinated dogs in New South Wales including Sydney (18), in 2011 there was a suspected outbreak in Victoria, although there is no report in the literature (11), and a case study reports distemper in ferrets in a rescue facility in Victoria (36). Wylie et al (2016) presented the most comprehensive contemporary assessment of CDV in Australia, reporting a total of 48 cases spread between 2006-2014. This study demonstrated the presence of CDV predominantly in the south-eastern states of Australia and one detection in the Northern Territory. Unfortunately, due to the disparity of the data it is unclear whether CDV was truly restricted to these areas, or, rather if CDV is widespread but no samples have been collected or outbreaks reported in other regions of Australia. Furthermore, due to the limited data it is unclear what the prevalence of CDV is in eastern Australia, in dogs or in other susceptible species. This is compounded by a lack of studies in wildlife in Australia such that it is unclear if CDV is maintained in dogs in Australia, or is maintained in wildlife such as Dingos (*Canis lupus dingo*), or introduced by feral animals such as foxes (*Vulpes vulpes*) (37) or ferrets (38). Due to these uncertainties, the origin and extent of the outbreak reported here remains opaque. While we report outbreaks of CDV at two research centres, we recognize that the outbreaks in these facilities may act as potential indicators for a larger outbreak within the Australian ferret population. From the reported outbreak in these facilities, we can only guess at which animals played a role as reservoirs in Victoria and were the source of the outbreak, the mechanism of transmission to ferret breeders and the extent of the outbreak in breeding facilities or elsewhere in the state.

Due to the complete lack of sequence data and only sporadic outbreaks and detection of CDV in Australia in the last 40 years, it is unclear whether previous CDV outbreaks in Australia were due to reversions of the vaccine strain or whether another lineage of CDV has been circulating on the continent. Through a combination of a discriminatory PCR assay and full genome sequencing we reveal a novel and highly divergent lineage of CDV infecting Australian ferrets. While this novel lineage is related to the “Vaccine/Asia-3/North America-1” lineages, a time-structured Bayesian phylogenetic analysis suggested that the novel field strain revealed here diverged from all other reported lineages well over 200 years ago. While we have identified a novel lineage, it is entirely unclear whether this constitutes an Australian lineage that has been geographically isolated since the divergence from the Vaccine/Asia-3/North America-1, or constitutes a lineage that was cryptically circulating elsewhere and has only recently been introduced to Australia. It is notable that there are a number of lineages circulating on most continents and in countries where CDV has been investigated (17, 39, 40), so it is not unreasonable to assume long - term circulation in Australia, and, whether additional viral diversity will be found in Australia remains to be determined.

In addition to detecting a novel lineage, we found that 51% of ferrets tested with a discriminatory PCR were still positive for the vaccine strain for up to 3 months after vaccination. The vaccine used in this study was a modified-live attenuated vaccine; the virus still has the ability to replicate in the animal and may have sufficient residual virulence to cause disease (41). Importantly, the safety and effectiveness of CDV vaccines are often tested in dogs, not mustelids (42). While the Protech C3 Vaccine is frequently used in Australia at present, there are other CDV vaccines on the global market that are subunit vaccines recommended specifically for use in ferrets, for example the Purevax Ferret Distemper Vaccine (Boehringer Ingelheim), which is a recombinant canarypox vector expressing the HA and F glycoproteins of canine distemper virus) that may prevent unwanted CDV symptoms seen in some ferrets (43). Despite some disease signs associated with the live attenuated CDV vaccine used in this study in ferrets, within 2 months of initiating vaccination the prevalence of the field strain dropped to zero and the ferret CDV outbreaks were curbed at both the Doherty Institute and ADCP. Whether this coincides with the end of the CDV outbreak in ferret breeding facilities or was linked to the introduction of vaccination is unclear as no epidemiological data from the breeding facilities was available. Genetic analysis demonstrated a large genetic difference between this putative Australian lineage and the vaccine used in this outbreak (Protech C3 Vaccine), with only ∼92% nucleotide similarity and 91% by amino acid similarity. Whether this genetic difference corresponds to an antigenic difference that would lead to low vaccine protection (44) is unclear but warrants further investigation. It is notable that there have been previously reported cases in both the Americas and in the UK of vaccine escape (23, 45), particularly when the Rockborn strain of the virus was used for vaccination (45).

Ferrets with signs of CDV in this outbreak varied in disease severity, in contrast to the very high mortality rates, with some outbreaks comprising a 100% mortality rate, reported previously in mustelids (11, 46, 47). This lower mortality, specifically at the Doherty Institute, may have been in part due to intervention measures that were used on suspected CDV infected ferrets, such as injections of vitamin A (36, 48), the brief time period between identification of disease signs and interventions (1-2 days), and the decision to cull ferrets at an early disease stage in order to control the spread of the outbreak. The disease signs observed in animals that required intervention did not display the full extent of morbidity previously reported (12, 13), however, this may similarly be confounded by the decision to intervene early in the disease course in order to contain the outbreak. Therefore, in the current investigation, the true pathogenicity of this new CDV strain remains unclear. A dedicated experimental challenge trial in ferrets will be required in order to determine this. Nevertheless, in ferrets submitted for histopathology, the typical profile of distemper pathology and multi-systemic virus replication were observed which suggests that, at least for a subset of infected ferrets, the highly divergent CDV strain described in our current study is capable of producing typical CDV-induced disease.

Overall CDV continues to pose a large disease burden, globally (49). In addition to dogs, this virus also has an impact on wildlife, with a number of examples of CDV infection in large cats, hyenas and jackals from many parts of Africa (8, 50). Due to the broad host range of CDV, cross-species infection is known to occur (50, 51), with the same lineages of CDV having been detected in canids, felids, mustelids and even seals. This is of concern, particularly in Australia, where several species of native fauna are currently at risk of extinction due to other factors (52). If this strain of CDV was to potentially infect these at-risk and immunologically naive animals, it could potentially result in widespread disease and high rates of mortality (6). One of the limiting factors in mounting an appropriate response would be the lack of established methods for CDV surveillance across the country (11). Specifically, following vaccination with a live attenuated vaccine, current diagnostic assays may not be able to accurately discriminate between the presence of vaccine or a circulating strain. Due to the geographic isolation of some parts of Australia it is possible that other genetically distinct strains of CDV are present in the country, such that the discriminatory PCR developed here may have limited value in an outbreak with a different lineage. Currently CDV diagnosis is still best achieved through a combination of histopathology and qRT-PCR and should be interpreted in the context of case history and clinical presentation (11). The detection of this novel lineage signals the need, not only for more widespread surveillance of CDV in Australia, but also for sequencing of any viruses that are detected to better understand the diversity of this lineage and to reveal any other lineages that may be circulating on the continent. As the ferrets reported in this study were sourced from regional breeders, the presence of CDV in this population may indicate that the virus is present in both domestic and wild animals in the state of Victoria. Our study describes the endpoint of an outbreak of CDV and highlights the crucial need for CDV surveillance and sequencing to better disentangle CDV ecology and evolution which is a key component for future outbreak response and control.

## Acknowledgements

We would like to thank the staff at the Biological Resource Facility at the Doherty Institute as well as the CSIRO Werribee Animal Health Facility for their assistance with animal husbandry. We would also acknowledge Mr Honglei Chen (ACDP) for technical assistance with NGS and Dr. Mark Ford for assistance with veterinary queries (ACDP). We acknowledge the use of the CSIRO Australian Centre for Disease Preparedness, grid.413322.5 in undertaking this research.

